# Human CD34^+^ progenitor-derived DC2 and DC3 subsets support CD16G-mediated HIV-1 uptake

**DOI:** 10.64898/2026.09.21.753077

**Authors:** Fernando Laguía, Patricia Pinol, Molly A. Vickers, Jon Izquierdo-Pujol, César Muñoz-Fontela, Javier Martinez-Picado, Patricia Resa-Infante

## Abstract

Dendritic cells (DCs) are central orchestrators of antiviral immunity but can be exploited by HIV-1 to facilitate systemic dissemination. HIV-1 recognition by CD169 triggers massive endocytosis (MEND)-related internalization of viral particles into sac-like compartments in monocyte-derived dendritic cells (MDDCs), thereby promoting *trans*-infection of CD4^+^ T cells. However, MDDCS represent only one DC subpopulation, and whether other physiologically relevant DC subsets exhibit similar behavior remains unclear. Moreover, endogenous CD169 expression and viral capture have not been fully characterized in cDC1, DC2, or pDCs, while remain unknown in DC3. We therefore investigated the contribution of these DC subsets to CD169-mediated HIV-1 capture and subsequent viral dissemination.

We used flow cytometry to assess CD169 expression and viral capture across DC subsets differentiated from CD34^+^ hematopoietic stem and progenitor cells (HSPC) obtained from bone marrow (BM) and cord blood (CB), two sources that differ in cellular maturity. Confocal microscopy was used to evaluate sac-like compartment formation following immune activation with IFN-α or LPS and treatment with MEND inhibitors. *Trans*-infection capacity was quantified by measuring luciferase activity in TZM-bl reporter cells.

We identified cDC1, DC2, DC3, moDC (similar to MDDCs), and pDC subsets following the differentiation of CD34^+^ HSPC from both CB and BM. Among these populations, DC2, DC3, and moDC displayed high CD169 expression, which was significantly upregulated by IFN-α or LPS stimulation. BM-derived DCs exhibited higher basal CD169 expression than CB-derived DCs, whereas stimulation induced greater CD169 upregulation in CB-derived DCs. DC2, DC3, and moDC efficiently captured HIV-1 particles in cultures derived from both sources, and viral uptake was further enhanced by immune activation. Viral internalization into sac-like compartments depended on CD169 binding and MEND-related mechanisms, ultimately facilitating HIV-1 transfer to target cells.

DC2 and DC3 are key dendritic cell subsets that support CD169-dependent HIV-1 capture and are associated with MEND-driven internalization into sac-like compartments and subsequent viral transfer to target cells, as was previously described for moDCs. We further observed distinct basal and inducible CD169 expression profiles in BM- and CB-derived DCs, consistent with source-dependent maturation states. These results identify DC2 and DC3 as key cellular contributors to HIV-1 dissemination.

## 1. Introduction

Myeloid antigen-presenting cells, including monocytes, macrophages, and dendritic cells (DCs), are key immune sentinels that detect pathogens and initiate immune responses. However, highly pathogenic enveloped viruses can exploit these protective cellular pathways to promote dissemination within the host (Perez-Zsolt et al., 2020). In particular, HIV-1 engages the CD169/Siglec-1 attachment receptor on activated myeloid cells, specifically monocyte-derived DCs (moDCs) and macrophages (Hammonds et al., 2017; Izquierdo-Useros et al., 2012, 2014; Pino et al., 2015; Puryear et al., 2012). CD169 recognizes viral membrane gangliosides and promotes virion accumulation in specialized vesicular structures referred to as sac-like compartments in DCs or virus-containing compartments in macrophages (Izquierdo-Useros et al., 2007, 2011, 2012; Pino et al., 2015; Puryear et al., 2012, 2013). From these structures, infectious particles can be transferred in *trans* to CD4^+^ T cells and disseminate through lymphoid tissues (Izquierdo-Useros et al., 2007; Pino et al., 2015; Wang et al., 2007). Similar CD169- dependent mechanisms have also been described for other enveloped viruses, including Ebola virus (EBOV) and severe acute respiratory syndrome coronavirus 2 (SARS-CoV-2) (Perez-Zsolt, Erkizia, et al., 2019; Perez-Zsolt et al., 2021), suggesting that sac-like compartments may represent a conserved myeloid cell-mediated pathway for viral dissemination. We recently demonstrated that massive endocytosis (MEND)-related mechanisms are triggered upon CD169-mediated viral attachment and contribute to HIV-1 internalization in monocyte-derived dendritic cells (MDDCs) generated *in vitro* from peripheral blood (Laguía et al., 2025). Because this model represents a specific inflammatory DC subset derived from primary monocytes (Coillard & Segura, 2019; Coutant et al., 2021; Sallusto & Lanzavecchia, 1994; Segura, 2022, 2025), it remains unknown whether CD169-mediated internalization operates similarly in other physiologically relevant DC subpopulations.

The limited availability of suitable DC experimental models has constrained functional studies of human DC biology (Han et al., 2019; Luo & Dalod, 2020). Primary DCs isolated from blood or tissues provide the closest physiological representation, but their low abundance limits experimental yield and robustness (Lubin et al., 2024). Consequently, MDDCs have been widely used because they can be generated in large numbers for mechanistic studies, although they primarily resemble inflammatory DC-like cells rather than steady-state DC subsets found *in vivo* (Collin & Bigley, 2018; Lundberg et al., 2013; Segura, 2022). In contrast, FLT3 ligand-driven differentiation of human CD34^+^ hematopoietic stem and progenitor cells (HSPC) from bone marrow (BM) or cord blood (CB) can generate multiple *bona fide* DC subsets, including cDC1, DC2, DC3, pDC, and provide a more physiologically relevant *in vitro* system that closely mirrors *in vivo* DC populations in gene expression and function (Anselmi et al., 2020; Balan et al., 2018; Lutz et al., 2023).

Although CD169-mediated HIV-1 uptake and *trans*-infection have been extensively characterized in the MDDC model, much less is known about HIV-1 capture and entry in other physiologically relevant myeloid populations, either *in vitro* or *in vivo* (Buffa et al., 2025). Available evidence indicates that CD169 expression is generally absent from cDC1, DC2, and pDCs (Buffa et al., 2025; Rhodes et al., 2021), whereas inflammatory signals can induce moderate CD169 transcriptional expression in DC3 *in vivo* (Dutertre et al., 2019; Winheim et al., 2021). Interestingly, DC2 cells have shown a moderate capacity to capture HIV-1 in *ex vivo* models under steady-state conditions, whereas cDC1 and pDC do not appear to support *trans*-infection, and experimental evidence regarding HIV-1 capture by DC3 remains limited (Buffa et al., 2025; Rhodes et al., 2021). However, most of these studies used *ex vivo* or tissue-derived cells, limiting controlled comparisons across DC subsets (Bertram et al., 2019; Buffa et al., 2025; Perez-Zsolt, Cantero-Pérez, et al., 2019; Rhodes et al., 2021; Rodriguez-Garcia et al., 2017). Moreover, CD169 expression under immune-activated conditions has not been systematically assessed in controlled *in vitro* systems such as CD34^+^ HSPC-derived DC cultures, which can generate multiple DC subsets and provide a useful platform for comparative analyses of viral uptake by myeloid cells.

Here, we used previously established *in vitro* differentiation protocols to generate cDC1, DC2, DC3, pDCs, and moDCs, the latter analogous to peripheral blood-derived MDDCs, from BM- or CB-derived CD34^+^ progenitors.. With this physiologically relevant system, we examined how immune activation modulates CD169 expression across DC subsets and whether CD169 induction is associated with enhanced HIV-1 particle capture. By comparing distinct DC populations under controlled experimental conditions, we sought to define the contribution of conventional DC subsets, particularly DC2 and DC3, to CD169-dependent viral uptake and to their potential role in myeloid cell-mediated HIV-1 dissemination.

## 2. Material and methods

### Cellular models

Human non-therapeutic grade cord blood HSPC from 2 different donor mixes (cat. no. 70008.3, Stem Cell); and HSPC from 2 individual bone marrow donors (cat. no. 70002, Stem Cell) were expanded and differentiated based on previous protocols (Balan et al., 2018). Thus, HSPCs were thawed and expanded for 7 days in 96-well round bottom plates at 40,000 cells/well/200 µL in Alpha Minimum Essential Medium without ribonucleosides and deoxyribonucleosides and supplemented with Glutamax (cat. no. 32561-029, Gibco), 10% FBS (cat. no. A5256701, Gibco), penicillin/streptomycin at 100 IU/ml (cat. no. 15070-022, Life Technologies), 2 mM Glutamine (cat. no. 25030-081, Gibco), 1mM sodium pyruvate (cat. no. 11360-070, Gibco), 50 µM β-mercaptoethanol (cat. no. 31350-010, Gibco), 25ng/ml hFLT3L (cat. no. 300-19, Peprotech), 5 ng/ml hIL7 (cat. no. 300-19, Peprotech), 5 ng/ml hTPO (cat. no. 300-18, Peprotech), and 2.5 ng/ml hSCF (cat. no. 300-07, Peprotech). Cells were harvested, counted and either immediately overlaid onto differentiation plates or cryopreserved in 90% FBS and 10% DMSO (cat. no. 276855, Merck) and stored in liquid nitrogen.

OP9 cell line (cat. no. CRL-2749, ATCC) was cultured at 37°C and 5% CO2 in feeder media that consist of Alpha Minimum Essential Medium without ribonucleosides and deoxyribonucleosides and supplemented with Glutamax (cat. no. 32561-029, Gibco), 20% FBS (cat. no. A5256701, Gibco), penicillin/streptomycin at 100 IU/ml (cat. no. 15070-022, Life Technologies), 2 mM Glutamine (cat. no. 25030-081, Gibco), 1mM sodium pyruvate (cat. no. 11360-070, Gibco) and 50 µM β-mercaptoethanol (cat. no. 31350-010, Gibco). For differentiation plates, OP9 stromal cells were seeded in a flat-bottom 24-well plate (12,500 cells/well/500μl) and expanded HSPC-derived cells were then overlayed (20,000 cells/mL/well) in feeder media supplemented with 15 ng/ml hFLT3L, 5 ng/ml hIL7, 5 ng/ml hTPO, and 2.5 ng/ml GM-CSF (cat. no. 300-03, Peprotech). Cells were incubated for 14 days at 5% CO2, 37°C. On day 7, half of the media was removed and replaced with double the initial concentration of cytokine.

HSPC-derived differentiated cells were stimulated for 48 h with IFN-α at 1000U/ml (cat. no. 11200-1, PBL assay science) or lipopolysaccharide at 100ng/ml (LPS, cat. no. L4391-1MG, Sigma-Aldrich).

TZM-bl cell line (obtained through the US National Institutes of Health [NIH] AIDS Research and Reference Reagent Program) was cultured DMEM (cat. no. 41966–029, Gibco) with 10% FBS and penicillin/streptomycin at 100 IU/ml. This cell line is an engineered HeLa-derived cell line that express stably CD4, CCR5 and CXCR4 viral receptors, and express luciferase reporter gene under an HIV-1 long terminal repeat (LTR).

### Immunophenotyping and virus binding by flow cytometry

The immunophenotype of expanded HSPC was evaluated by conventional flow cytometry with cell surface markers indicated in **Table 1**. Cells were blocked with human immunoglobulins (cat. No. 44206, Privigen, Behring CSL) for 20 min at room temperature before staining with indicated antibodies for 30 min at 4°C. Cells were then washed and fixed with 4% paraformaldehyde 4% before acquisition in BD FACSCanto II flow cytometer (BD Biosciences, San Jose, CA, USA). Resulting data was analyzed with FlowJo v10 (BD Biosciences, Ashland, OR, USA). Each single marker expression was analyzed over the total live cell population.

**Table 1.** List of antibodies used in conventional flow cytometry.

| Marker (clone) | Fluorophore | Reference | Provider | Dilution |
| --- | --- | --- | --- | --- |
| CD3 (SK7) | AmCyan | 339186 | BD | 1:20 |
| CD25 (ML5) | PE-Cy7 | 561646 | BD | 1:20 |
| CD4 (SK3) | PerCP-Cy5.5 | 345770 | BD | 1:20 |
| CD34 (8G12) | PE | 340669 | BD | 1:10 |
| CD14 (MφP9) | FITC | 345784 | BD | 1:20 |
| CD8 (SK1) | APC-Cy7 | 561945 | BD | 1:40 |
| CD69 (FN50) | APC | 555533 | BD | 1:5 |
| Live/Dead | Blue | L23105 | Thermofisher | 1:500 |

The immunophenotype of differentiated HSPC was evaluated by spectral flow cytometry. HSPC-derived differentiated cells were transferred to U-Bottom 96-well polypropylene plates (cat. no. 267245, ThermoFisher), incubated for 10 min with TruStain FcX to block Fc receptors (cat. no. 422301, Biolegend). Cells were then stained with the Zombie NIR viability Kit (cat. no. 423105, Biolegend) for 10 min, washed and stained for 20 min with conjugated antibodies to measure cell surface markers indicated in **Table 2**. Finally, cells were washed and fixed with cytofix buffer (cat. no. 554655, BD) for 15 min and acquired on Cytek Aurora spectral flow cytometer (Cytek Biosciences, Fremont, CA, USA). Resulting data was analyzed with FlowJo (BD Biosciences, Ashland, OR, USA) and OMIQ (Dotmatics, Boston, MA, USA).

**Table 2.** List of antibodies used in spectral flow cytometry.

| Marker (clone) | Fluorophore | Reference | Provider | Dilution |
| --- | --- | --- | --- | --- |
| CD14 (63D3) | SB550 | 367148 | Biolegend | 1:80 |
| CD141 (1A4) | BV711 | 563155 | BD | 1:200 |
| CD11c (B-ly6a) | BV650 | 563404 | BD | 1:200 |
| CD206 (19.2) | PE-Cy7 | 25-2069-42 | Thermofisher | 1:200 |
| CD123 (6H6) | PerCP-Cyanine5.5 | 45-1239-42 | Thermofisher | 1:200 |
| CLEC9A (8F9) | PE | 353804 | Biolegend | 1:200 |
| CLEC10A (H037G3) | APC | 354706 | Biolegend | 1:200 |
| CD1c (L161) | BV421 | 331526 | Biolegend | 1:66 |
| CD169 7-239 | R718 | 751877 | Biolegend | 1:100 |
| Live/Dead | Zombie NIR | 423105 | Biolegend | 1:500 |

As detailed in gating strategy (**Fig. 1B**), moDCs were defined as CD14^+^, CD206^+^, CD11c^+^, CD1c^-^. DC3 were defined as CD14^+^, CD206^+^, CD11c^+^, CD1c^+^. cDC1 were defined as CD14^-^, CD123^-^, CD141^+^ and CLEC9A^+^. DC2 were defined as CD14^-^, CD123^-^, CD141^-^, CLEC9A^-^, CD206^+^, CD11c^+^, CD1c^+^ and CLEC10A^+^. pDCs were defined as CD14^-^, CD123^+^. For each subset, CD169 median fluorescence intensity (MFI) was measured at different activation conditions. FlowSOM algorithm was used to identify and annotate immune population clusters. Only clusters above 1000 events were considered for further analysis. Then, the same markers were used for dimensionality reduction with uniform manifold approximation and projection (UMAP) for clustering.

**Figure 1.**
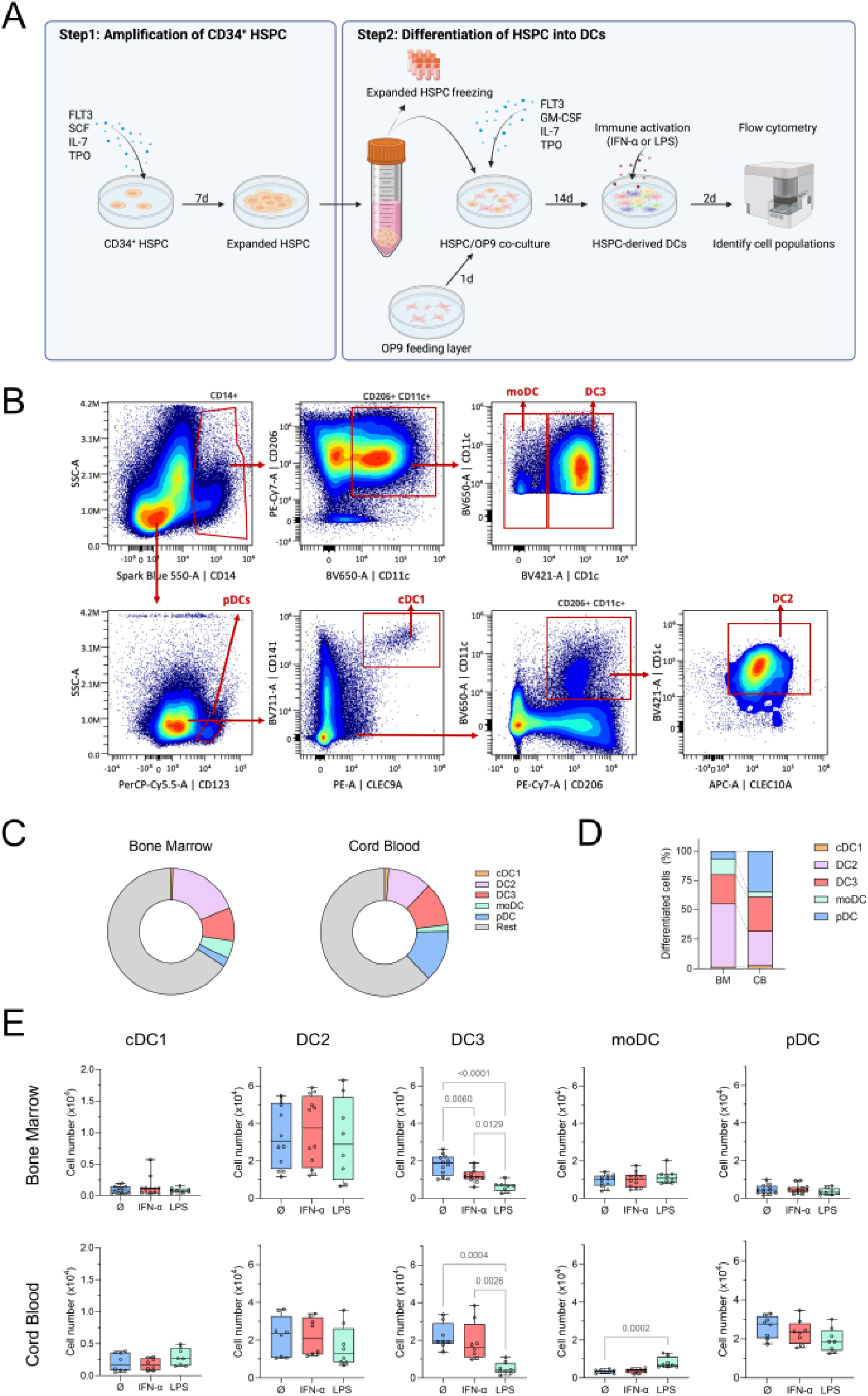
CD34^+^ HSPCs from bone marrow and cord blood generate multiple dendritic cell subsets *in vitro*. CD34^+^ HSPC from BM or CB were expanded and differentiated into DCs under FLT3L-based culture conditions. Differentiated cells were left untreated or stimulated with IFN-α or LPS for 48 h before analysis. **(A)** Schematic representation of the expansion and differentiation protocol used to generate HSPC-derived DCs from BM and CB progenitors. **(B)** Flow cytometry gating strategy used to identify cDC1, DC2, DC3, moDC, and pDC subsets within HSPC-derived cultures. **(C)** Frequency of DC subsets expressed as percentage of total live cells in BM- and CB-derived cultures. **(D)** Frequency of each DC subset expressed as a percentage of total differentiated DCs. **(E)** Absolute numbers of each DC subset after stimulation with IFN-α or LPS. Cell counts were normalized to 200,000 live cells. Dots represent individual experiments, and boxes indicate the median and interquartile range. Statistical significance was assessed using one-way ANOVA. Significant *P* values for each activation condition are shown in the graph.

When indicated, HSPC-derived cells were pulsed for 2 h with saturating levels of VLP_HIV_ generated from the plasmid pHIV-Gag-eGFP as previously described (Laguía et al., 2025) to assess VLP_HIV_ binding by spectral flow cytometry. These samples were not considered for CD169^+^ MFI analyses, as CD169 receptor occupancy by VLP_HIV_ interferes in anti-CD169 antibody binding.

### Cell imaging by confocal microscopy

To study VLP_HIV_ distribution under steady-state and activation conditions, cells were pulsed for 6 h with saturating levels of VLP_HIV_. PBS-washed cells were attached to glass coverslips coated with poly-L-lysine solution (cat. no. P8920, Merck). Then, cells were fixed for 20 min at 4°C in 4% paraformaldehyde (cat. no. P6148, Merck) before storage in PBS containing 0.5% w/v BSA (cat. no. A966-506, Sigma-Aldrich).

To evaluate the effect of MEND inhibition on VLP_HIV_ distribution, IFN-α–activated cells were treated for 2 h at 37°C with either 100 nM wortmannin (PI3K inhibitor; cat. no. PHZ1301, ThermoFisher) or 25 µM 2- bromopalmitate (2-BP; palmitoyl-acyl transferase inhibitor; cat. no. 238422, Merck) prior to a 3-h VLP_HIV_ pulse. After VLP_HIV_ incubation, cells were washed and transferred to glass coverslips as described.

Fixed HSPC-derived cells were treated for 1 h with a permeabilization buffer containing 0.1% saponin and 0.5% BSA in PBS (1 mg/ml saponin, cat. No. SAE0073, Merck; 5 mg/ml BSA, cat. no. A7906 Merck). Subsequently, cells were incubated for 1 h with 20 ng/ml of anti-CD169 antibody clone #6H9 (Perez-Zsolt, Erkizia, et al., 2019) conjugated with Abberior STAR RED-NHS dye (cat. no. STRED-0002-1MG, Abberior GmbH) and anti-CD1c polyclonal antibody (cat. no. PA5117456, ThermoFisher). Then, cells were washed and stained for 1h with an anti-rabbit secondary antibody labelled with AF568 (cat. no. A11011, ThermoFisher). Nuclei were stained with 300 nM DAPI solution in PBS (cat. no. D1306, ThermoFisher) for 5 min. Coverslips were mounted onto glass slides with ProLong Glass Antifade mounting media (cat. no. P36982, ThermoFisher) and sealed with nail polish. Fixed samples were then imaged using Andor Dragonfly 505 spinning disk confocal microscope.

To evaluate viral particle distribution in cells, we first determined cell boundaries based on anti-CD1c antibody signal and selected VLP_HIV_^+^ based on eGFP signal using Fiji/ImageJ software. Then, we utilized a script based on the circularity of the eGFP channel to determine whether cells were polarizing VLP_HIV_ (circularity value = 0 - 0.7) or whether cells formed sac-like compartment (circularity value = 0.7 - 1) as previously described (Laguía et al., 2025). We set a minimum sac-like compartment size of 1 micro and other VLP_HIV_^+^ cells with eGFP signal smaller than this were assigned to random binding category.

### HIV-1 *trans*-infection assay

IFN-α treated HSPC-derived DCs were pulsed with HIV-1_NL4–3_ virus with an MOI of 0.1 for 24h. For blocking experiments, cells were pre-incubated for 15 min at room temperature with either anti-CD169 antibody (clone 7-239, cat. no. ab199401, Abcam) or an isotype-matched control antibody (mouse IgG, cat. no. 554721, BD) prior to HIV-1 exposure. After extensive washing with cold PBS, cells were co-cultured with the reporter cell line TZM-bl at different ratios of HSPC-derived DCs vs. TZM-bl (4:1, 1:1 or 1:4), while maintaining a constant number of TZM-bl cells per well. Cells were assayed for luciferase activity 48 h later (BrightGlo luciferase system; Promega) in a EnSight™ Multimode Plate Reader (Perkin Elmer). Background values obtained from TZM-bl monocultures were measured for each experiment to correct background signal in experimental wells.

### Statistical analysis

Two independent experiments were performed using CB-derived HSPCs from each cord blood donor- mix and three independent experiments were performed using BM-derived HSPCs from each bone marrow donor to evaluate cell subset distribution, CD169 expression and VLP_HIV_ binding. Moreover, cell immunophenotyping and CD169 positivity rate were quantified in duplicate technical replicates. One-way ANOVA test was used to compare cell frequencies and two-way ANOVA was used to compare MFI measurements among cellular conditions. In addition, for *trans*-infection assays, five and three independent experiments were performed using for CB-derived and BM-derived HSPCs, respectively, and included duplicate technical replicates. Unpaired t-test was used to compare normalized values and one-way ANOVA test was used to compare absolute values. All statistical analyses were performed using GraphPad Prism 10 (GraphPad Software). Differences were considered statistically significant for p values < 0.05.

### Ethics and biosafety statements

The study was approved by the institutional review board for biomedical research of Hospital Germans Trias i Pujol (Ref. PI-24-201).

## 3. Results

### 3.1. CD34^+^ HSPC progenitors from bone marrow and cord blood generate multiple dendritic cell subsets for functional studies

Given that the MDDC model represents a specific inflammatory DC population and that primary DC availability is limited, we differentiated CD34^+^ HSPCs from BM and CB into DCs using cytokine-based culture conditions (**Fig. 1A**). Cells were first cultured for 7 days with an expansion cytokine cocktail containing FLT3L, SCF, IL-7 and TPO, followed by 14 days of culture with a differentiation cytokine cocktail containing FLT3L, GM-CSF, IL-7 and TPO.

Both BM- and CB-derived HSPCs showed an approximately fivefold expansion after the 7-day expansion phase, together with a shift in CD34 expression, resulting in a mixed CD34^+^/CD34^−^ population indicative of loss of stemness and initiation of differentiation (**Fig. S1**). Interestingly, BM-derived HSPCs showed increased CD14 and CD4 expression (**Fig. S1D**), supporting progression toward myeloid differentiation during the expansion phase. BM-derived HSPCs also displayed a slight increase in CD25 expression, suggesting activation.

Following the 14-day differentiation process, the frequency of HSPC-derived DC subpopulations was analyzed by spectral flow cytometry (**Fig. 1B**). To validate manual gating, we further analyzed flow cytometry data using the unsupervised clustering algorithm FlowSOM. This analysis identified clusters corresponding to the manually gated populations, supporting our choice of selected markers used to define subsets in this study (**Fig. S2**). Approximately 30% of cells displayed phenotypic profiles consistent with differentiated DC subsets, including cDC1, DC2, DC3, moDCs, and pDCs, in cultures derived from both BM and CB HSPC (**Fig. 1C**). DC2 (18% in BM vs. 11% in CB), DC3 (9% in BM vs. 11% in CB), and moDCs (5% in BM vs. 2% in CB) were the most abundant subpopulations in both culture systems, whereas pDCs were enriched in CB-derived cultures compared with BM-derived cultures (2% in BM vs. 13% in CB) (**Fig. 1D**). To evaluate the impact of immune activation on subset composition, differentiated cells were stimulated with IFN-α or LPS for 48 h. Most subsets remained stable in absolute numbers compared to mock-treated conditions, except for the DC3 population, which decreased following IFN-α or LPS stimulation in cultures derived from both BM and CB (**Fig. 1E**).

Taken together, these results demonstrate that multiple DC subsets can be simultaneously generated from CD34^+^ progenitors *in vitro*, providing a suitable platform for downstream functional analyses across DC subsets. Due to their low frequency, the cDC1 subset was not included in subsequent analyses.

### 3.2. CD16G expression is subset-specific and inducible by immune activation in HSPC-derived DCs

To assess CD169 expression across HSPC-derived DC subsets, we analyzed its expression under steady-state conditions and following immune activation with IFN-α or LPS by spectral flow cytometry (**Fig. 2A**). At the level of total Dpopulations, BM-derived cultures displayed a higher proportion of CD169^+^ cells (∼80%) than CB-derived cultures under steady-state conditions (∼20%). IFN-α stimulation increased the proportion of CD169^+^ cells in both BM- and CB-derived cultures (∼90% and ∼60%, respectively). In contrast, LPS-induced CD169 upregulation was observed only in BM-derived cultures, reaching approximately 90% CD169^+^ cells (**Fig. 2B**).

**Figure 2.**
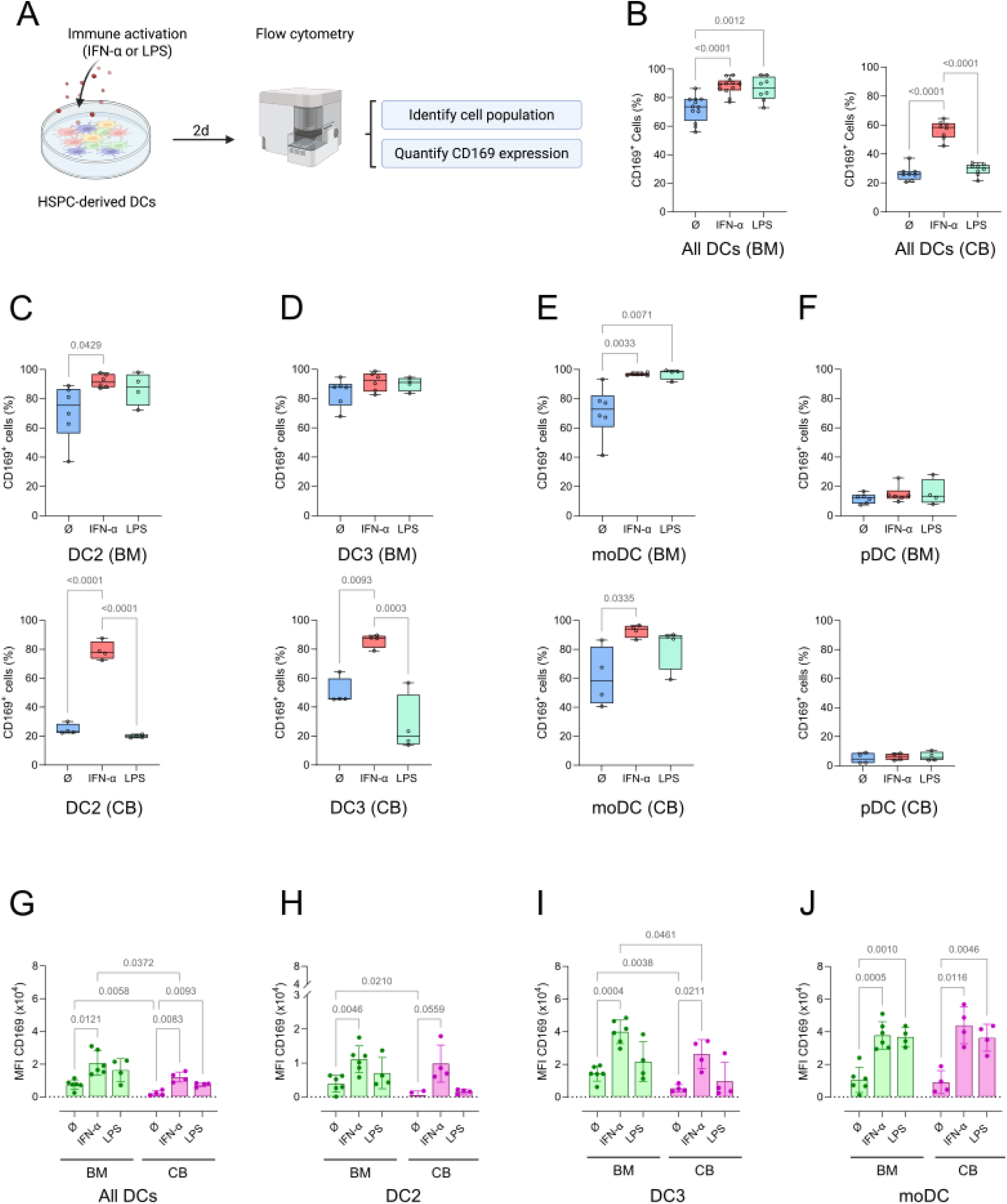
CD16G expression is differentially regulated across HSPC-derived DC subsets. BM- and CB-derived cells were left untreated or stimulated with IFN-α or LPS for 48 h and analyzed by spectral flow cytometry. **(A)** Experimental workflow used for CD169 expression analysis. **(B-F)** Percentage of CD169^+^ cells among total DCs **(B)**, DC2 **(C)**, DC3 **(D)**, moDCs **(E)** and pDCs **(F)** derived from BM and CB cultures. Dots represent individual experiments, and boxes indicate the median and interquartile range. Statistical significance was assessed using one-way ANOVA. Significant *P* values for each phenotype are shown in the graph. **(G-J)** Median fluorescence intensity (MFI) of CD169 staining in total DCs **(G)**, DC2 **(H)**, DC3 **(I)** and moDCs **(J)**. Dots represent individual experiments, and bars indicate the mean of MFIs. Statistical significance was assessed using two-way ANOVA. Significant *P* values for each phenotype are shown in the graph.

We next examined CD169 expression across individual DC subsets. Under steady-state conditions, the proportion of CD169^+^ DC2 cells was higher in BM-derived cultures (∼75%) than in CB-derived cultures (∼20%) (**Fig. 2C**). IFN-α stimulation markedly increased the proportion of CD169^+^ DC2 cells in both BM- and CB-derived cultures (∼90% and ∼80%, respectively). In contrast, LPS stimulation increased the proportion of CD169^+^ DC2 cells in BM-derived cultures (∼90%) but had minimal effect in CB-derived cultures (∼20%). DC3 exhibited a higher basal proportion of CD169^+^ cells than DC2, particularly in CB-derived cultures (90% in BM-derived DC3 and 45% in CB-derived DC3) (**Fig. 2D**). IFN-α further increased the proportion of CD169^+^ cells in both BM- and CB-derived DC3 cultures (∼90%), whereas LPS-induced upregulation was restricted to BM-derived cultures (90%) and was not observed in CB-derived cultures (20%). In contrast, moDCs displayed a consistently high proportion of CD169^+^ cells in both BM- and CB-derived cultures (∼70% and ∼60%, respectively), which further increased after IFN-α or LPS stimulation to more than 90% (**Fig. 2E**). pDCs exhibited low CD169 expression and did not show induction upon stimulation; therefore, they were excluded from subsequent analyses (**Fig. 2F**).

Analysis of CD169 median fluorescence intensity (MFI) revealed patterns consistent with the frequency data, indicating increases both in the proportion of CD169^+^ cells and in per-cell CD169 expression levels (**Fig. 2G**). LPS-induced increases in CD169 MFI were primarily observed in BM-derived cultures, with limited effects in CB-derived cultures. Across subsets, basal CD169 expression was lowest in DC2. Intermediate expression levels were observed in CB-derived DC3 and moDCs, whereas BM-derived DC3 and moDCs exhibited the highest expression levels (**Fig. 2H**). DC2 cells showed an approximately twofold increase in CD169 MFI upon IFN-α stimulation in both BM- and CB-derived cultures, whereas LPS had minimal effect. Similarly, DC3 cells showed an approximately twofold increase in CD169 expression in response to IFN-α across both sources, whereas LPS-induced upregulation was restricted to BM-derived cultures. In contrast, moDCs exhibited a strong increase in CD169 expression after both IFN-α and LPS stimulation, irrespective of progenitor source. Notably, CD169 expression levels were consistently higher in DC3 and moDCs than in DC2 across all activation conditions (**Fig. 2H**). Consistent with the flow cytometry data, confocal microscopy showed sustained CD169 expression in BM-derived DCs and inducible CD169 expression after IFN-α stimulation in CB-derived DCs (**Fig. S3**).

In summary, these data indicate that DC2, DC3, and moDC subsets express basal levels of CD169 that are differentially upregulated by immune activation in BM- and CB-derived cultures. These subset- and activation-dependent differences in CD169 expression prompted us to examine the capacity of these DC populations to bind HIV-1 particles.

### 3.3. HIV-1 particle uptake by dendritic cells correlates with CD16G expression across subsets

We next assessed the capacity of HSPC-derived DCs to capture HIV-1-like particles (VLP_HIV_). Cells were pulsed with VLP_HIV_ for 2 h, and viral binding was quantified by spectral flow cytometry (**Fig. 3A**). At the total DC population level, BM-derived cultures exhibited a higher proportion of VLP_HIV_^+^ cells (∼50%) than CB-derived cultures (∼15%) under steady-state conditions (**Fig. 3B**). IFN-α stimulation increased VLP_HIV_ binding in both BM- and CB-derived cultures, reaching approximately 80% and 70% VLP_HIV_^+^ cells, respectively. In contrast, LPS stimulation further increased VLP_HIV_ binding in BM-derived cultures (>80%), whereas a more modest increase was observed in CB-derived cultures (∼60%).

**Figure 3.**
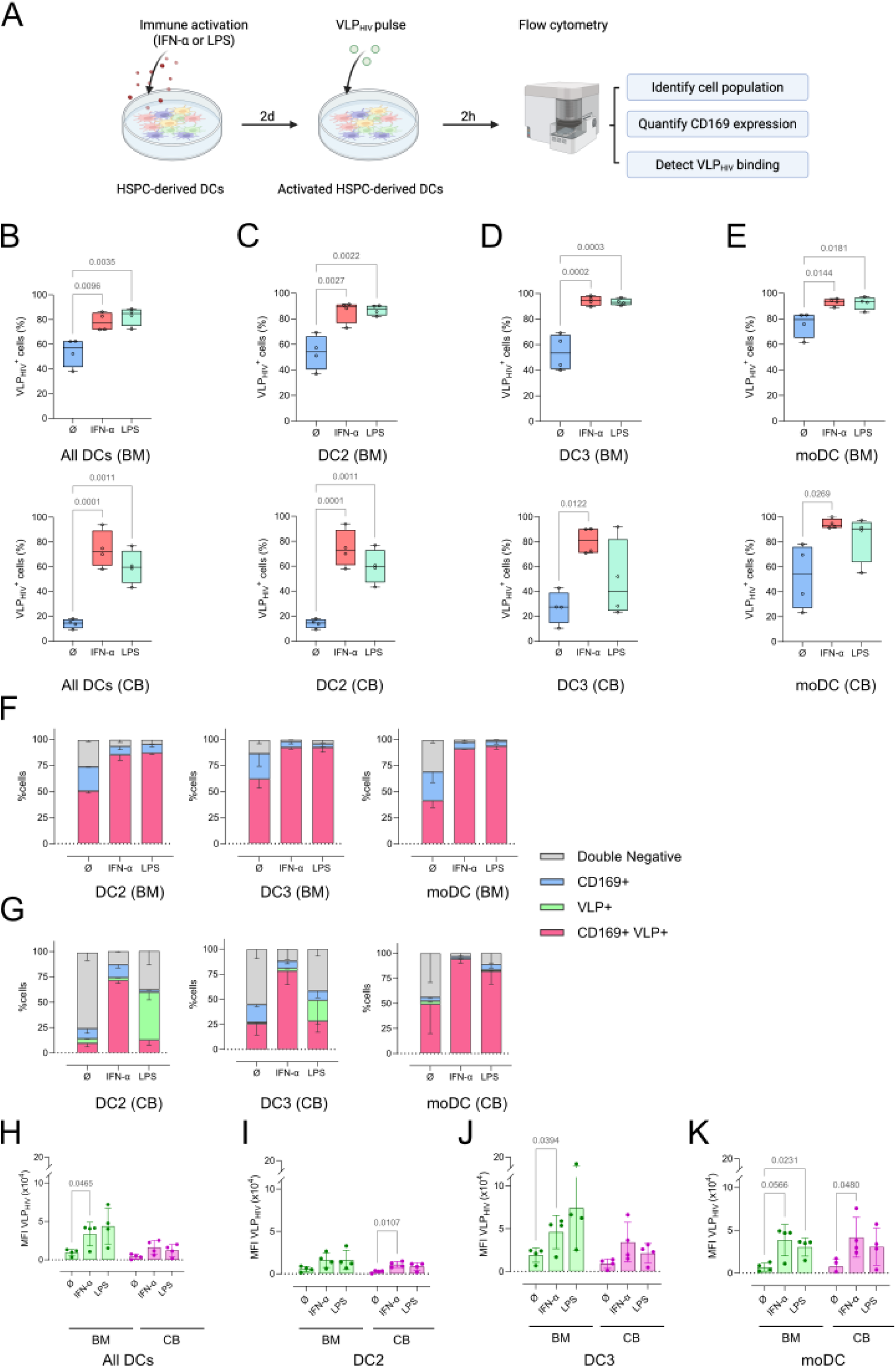
HIV-1 particle capture correlates with CD16G expression across HSPC-derived DC subsets. HSPC- derived DCs were left untreated or stimulated with IFN-α or LPS for 48 h before incubation with HIV-1 virus-like particles (VLPHIV). Dots represent individual experiments, and bars indicate mean values. Statistical significance was assessed using two-way ANOVA. **(A)** Experimental workflow used to assess VLPHIV binding. **(B-E)** Frequency of VLPHIV^+^ cells among total DCs **(B)**, DC2 **(C)**, DC3 **(D)**, and moDCs **(E)** derived from BM and CB cultures. Dots represent individual experiments, and boxes indicate the median and interquartile range. Statistical significance was assessed using one-way ANOVA. Significant *P* value for each phenotype is shown in the graph. **(F-G)** Distribution of VLPHIV and CD169 expression within DC subsets from BM-derived **(F)** and CB-derived **(G)** cultures. Bars represent the mean proportion of CD169^+^/VLPHIV^+^ double-positive cells, CD169 single-positive cells, VLPHIV single-positive cells, and double-negative cells. **(H-K)** Median fluorescence intensity (MFI) of GFP-labelled VLPHIV in total DCs **(H)**, DC2 **(I)**, DC3 **(J)**, and moDCs **(K)**. Dots represent individual experiments, and bars indicate the mean of MFIs. Statistical significance was assessed using two-way ANOVA. Significant *P* values for each phenotype are shown in the graph.

We then analyzed VLP_HIV_ binding across individual DC subsets. Under steady-state conditions, DC2 showed a higher proportion of VLP_HIV_^+^ cells in BM-derived cultures (∼50%) than in CB-derived cultures (∼20%) (**Fig. 3C**). IFN-α stimulation markedly increased binding in both sources (>80%), whereas the effect of LPS was more divergent, including a strong increase in BM-derived cultures (∼90%) but a more limited responses in CB-derived cultures (∼50%). DC3 followed a similar pattern (**Fig. 3D**). In BM-derived cultures, high basal binding (∼80%) increased further to near-saturation levels after IFN-α or LPS stimulation. In contrast, CB-derived DC3 displayed lower basal binding (∼25%), which increased substantially after IFN-α stimulation (∼80%), whereas LPS had minimal effect (**Fig. 3D**). In moDCs, VLP_HIV_ binding was comparable between BM- and CB-derived cultures (>60% at baseline) and increased further after IFN-α or LPS stimulation, reaching ∼90% in both sources (**Fig. 3E**).

To evaluate the association between CD169 expression and viral binding, we analyzed CD169 expression within the VLP_HIV_^+^ population. In BM-derived cultures, VLP_HIV_^+^ cells were consistently CD169^+^ across all subsets, indicating a strong association between CD169 expression and viral capture (**Fig. 3F**). Similarly, in CB-derived cultures, VLP_HIV_ binding was largely restricted to CD169^+^ cells under steady-state conditions and after IFN-α stimulation (**Fig. 3G**). However, following LPS stimulation, a substantial fraction of VLP_HIV_^+^ DC2 and DC3 cells in CB-derived cultures did not express CD169, suggesting that CD169-independent binding mechanisms may also contribute under these conditions.

Analysis of VLP_HIV_ MFI further supported these observations, showing increased signal intensity consistent with enhanced viral binding after IFN-α and LPS stimulation in total DC populations (**Fig. 3H**). At the subset level, IFN-α induced a marked increase in VLP_HIV_ capture in DC2 (**Fig. 3I**) and DC3 (**Fig. 3J**) across both BM- and CB-derived cultures, whereas LPS effects were more variable and generally limited in CB-derived subsets. In contrast, moDCs showed robust increases in viral capture under both activation conditions, reaching comparable levels irrespective of progenitor source (**Fig. 3K**).

In summary, HSPC-derived DCs efficiently capture VLP_HIV_, and this process is enhanced by immune activation in a subset-dependent manner that closely parallels CD169 expression patterns. These findings prompted us to investigate whether VLP_HIV_ capture leads to the formation of intracellular sac-like compartments.

### 3.4. HSPC–derived DCs form sac-like compartments upon HIV-1 binding via MEND-dependent mechanisms

To determine whether HSPC-derived DCs form sac-like compartments after viral binding, cells were stimulated with IFN-α or LPS and pulsed with VLP_HIV_ for 6 h to analyze viral particle distribution along the plasma membrane by confocal microscopy (**Fig. 4A**). Cells were classified into four phenotypic categories: cells negative for VLP_HIV_, random VLP_HIV_ distribution across the plasma membrane, VLP_HIV_ polarization at a discrete region of cell membrane, and VLP_HIV_ internalization into sac-like compartments. CD169 localization was assessed in parallel. Across all conditions, VLP_HIV_ colocalized with CD169, supporting an association between CD169 expression and sac-like compartment formation. In BM-derived DCs, a high proportion of cells displayed sac-like compartments after 6 h of VLP_HIV_ incubation, and this proportion further increased after IFN-α or LPS stimulation, with most cells showing sac-like compartment formation (**Fig. 4C**). In contrast, CB-derived DCs exhibited limited sac-like compartment formation under control conditions but a marked increase after IFN-α stimulation (**Fig. 4D**). LPS induced a more modest increase in sac-like compartment formation in CB-derived DCs, with many cells displaying a more dispersed VLP_HIV_ distribution at the cell surface.

**Figure 4.**
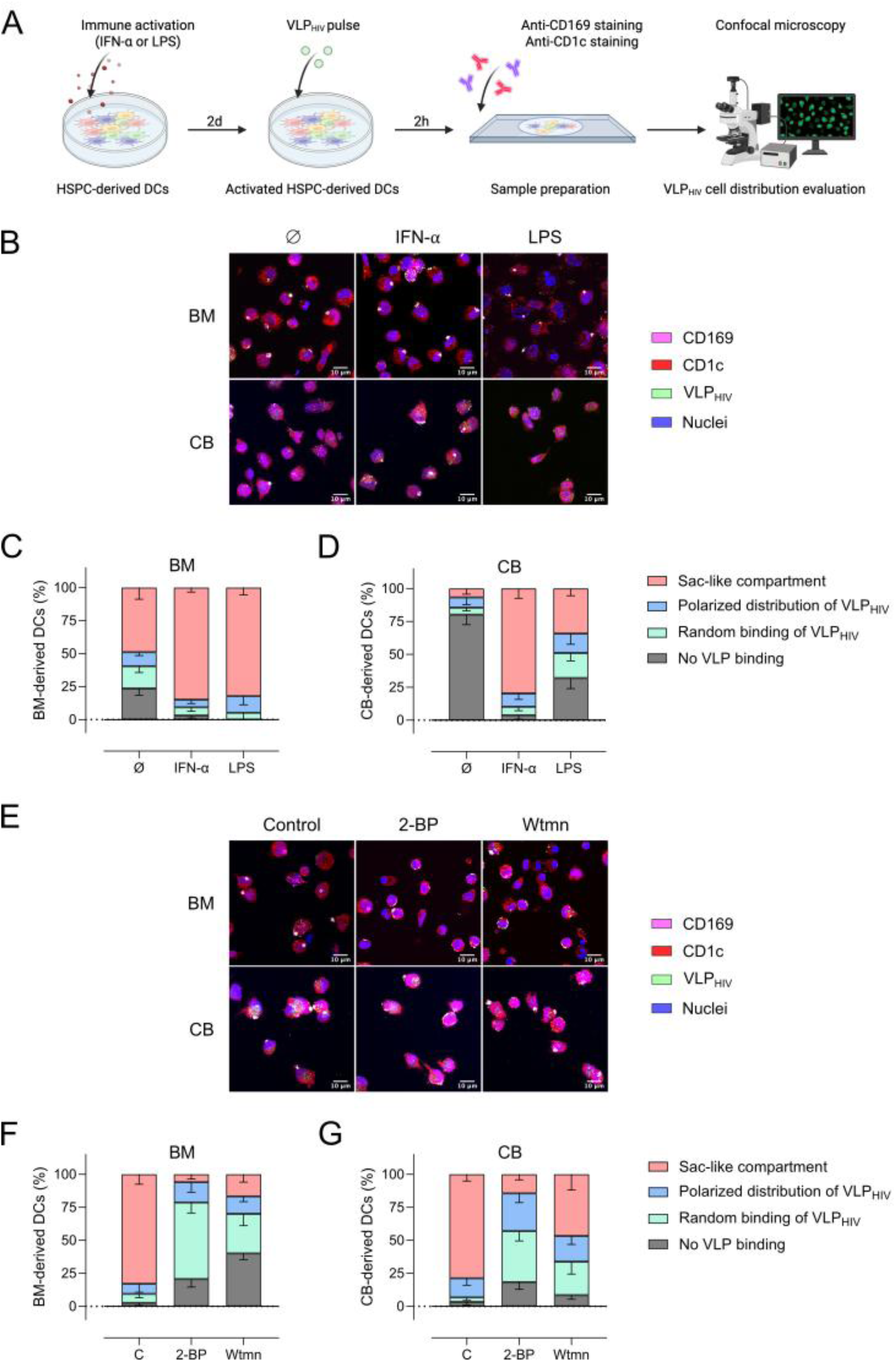
HSPC-derived DCs cells form sac-like compartments through MEND-dependent mechanisms following HIV-1 particle capture. BM- and CB-derived cultures were left untreated or stimulated with IFN-α or LPS before incubation with VLPHIV. Cells were fixed and stained for CD169 (magenta) and CD1c, used as a marker to delineate the cell volume (red). VLPHIV particles are shown in green and nuclei in blue. **(A)** Experimental workflow for confocal microscopy analysis of VLPHIV distribution. **(B)** Representative confocal images showing VLPHIV distribution in BM- and CB-derived cultures under the indicated activation conditions. **(C-D)** Quantitative analysis of VLPHIV distribution in BM-derived **(C)** and CB-derived **(D)** cultures. At least 100 cells were evaluated for each experimental condition. Cells were classified into four categories: no VLPHIV binding, random VLPHIV binding, polarized VLPHIV distribution, and sac-like compartment formation. Bars represent mean values from independent experiments, and error bars indicate the standard deviation. **(E)** Representative confocal images of IFN-α-stimulated BM- and CB-derived cultures either untreated (C) or treated with the MEND inhibitors 2-Bromopalmitate (2-BP) or wortmannin (Wtmn) before VLPHIV exposure. **(F-G)** Quantitative analysis of VLPHIV distribution in BM-derived **(F)** and CB-derived **(G)** cultures following inhibitor treatment. Cells were classified using the same criteria as in panels C–D. Bars represent mean values from independent experiments, and error bars indicate the standard deviation.

To assess the involvement of MEND in sac-like compartment formation, IFN-α–activated cells were pretreated with MEND inhibitors, specifically the palmitoylation inhibitor 2-BP or the PI3K inhibitor wortmannin as previously described. After treatment, HSPC-derived DCs were exposed to VLP_HIV_ for 3 h and viral particle distribution was assessed as described above (**Fig. 4E**). Both treatments markedly reduced sac-like compartment formation in BM- and CB-derived DCs (**Fig. 4F-G**). In particular, 2-BP strongly impaired sac-like compartment formation, whereas wortmannin reduced the number of sac-like compartments and also resulted in a more dispersed or polarized distribution of VLP_HIV_ along the plasma membrane.

Together, these results are consistent with the involvement of MEND-related mechanisms in sac-like compartment formation during HIV-1 particle capture in HSPC-derived DCs.

### 3.5. CD16G-mediated HIV-1 capture contributes to *trans*-infection of CD4^+^ T cells

To determine whether CD169-mediated viral capture contributes to functional HIV-1 *trans*-infection, IFN-α-activated BM- and CB-derived DCs were exposed to replication-competent HIV-1 in the presence or absence of a blocking anti-CD169 antibody, using an isotype-matched antibody as a control (**Fig. 5A**). After extensive washing, DCs were co-cultured with TZM-bl reporter cells at different DC:TZM-bl ratios (4:1, 1:1 and 1:4), and *trans*-infection was quantified by luciferase activity. In both BM- and CB-derived cultures, HIV-1-exposed DCs efficiently transferred infection to TZM-bl cells, whereas negligible reporter activity was detected in non-virus controls (**Fig. 5B**). The isotype control antibody did not substantially affect *trans*-infection levels, indicating that antibody addition itself did not interfere with viral transfer. Absolute luciferase values revealed higher reporter activity in TZM-bl cells co-cultured with CB-derived DCs than BM-derived DCs across conditions (**Fig. S4**).

**Figure 5.**
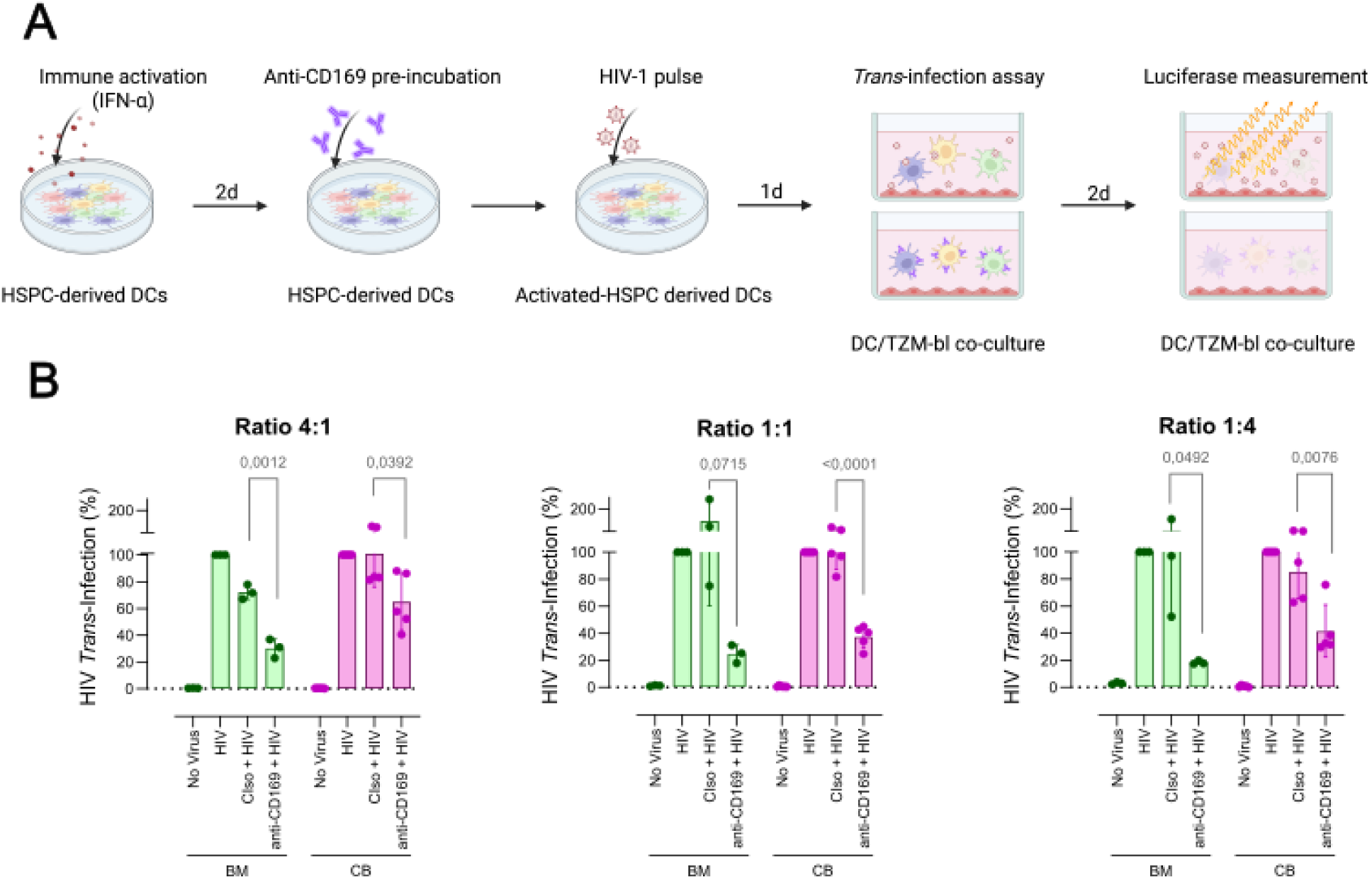
CD16G-mediated HIV-1 capture contributes to *trans*-infection of CD4^+^ target cells. IFN-α-activated HSPC-derived DCs from BM and CB cultures were exposed to replication-competent HIV-1 in the presence or absence of a blocking anti-CD169 antibody. An isotype-matched antibody was used as a control. After extensive washing, DCs were co-cultured with TZM-bl reporter cells, and HIV-1 *trans*-infection was quantified by luciferase activity. **(A)** Experimental workflow for the HIV-1 *trans*-infection assays. **(B)** Relative *trans*-infection efficiency of BM-derived and CB-derived DCs measured at DC:TZM-bl ratios of 4:1, 1:1 and 1:4. Values were normalized to the HIV-1 condition, which was set to 100%. Bars represent mean values from independent experiments, and error bars indicate the standard deviation. Statistical significance was assessed using unpaired t-test comparing isotype control and anti-CD169 conditions.

Blocking CD169 consistently reduced HIV-1 *trans*-infection in both BM- and CB-derived DCs cultures across all co-culture ratios. Relative to HIV-1-exposed controls, anti-CD169 treatment decreased *trans*- infection by approximately 70% in BM-derived co-cultures and by approximately 35%, 63% and 58% in CB-derived co-cultures at DC:TZM-bl ratios of 4:1, 1:1 and 1:4, respectively (**Fig. 5B**). Comparison of anti-CD169 and isotype control conditions demonstrated a significant reduction in trans-infection in all BM- and CB-derived cultures, except for BM-derived cultures at the 1:1 ratio. Analysis of absolute luciferase values showed a similar trend than relative values, although statistical significance was only reached in BM-derived cultures, likely reflecting the variability of absolute reporter signals and the limited number of biological replicates (**Fig. S4)**. These results indicate that CD169-mediated viral capture makes a major contribution to the *trans*-infection capacity of HSPC-derived DCs.

Taken together, these results demonstrate that CD169-mediated HIV-1 capture contributes substantially to the capacity of CD34^+^ HSPC-derived DCs to *trans*-infect CD4^+^ T cells. The partial inhibition observed after CD169 blockade further suggests that additional CD169-independent pathways may also participate in viral transfer.

## 4. Discussion

Developing robust and reproducible experimental models that accurately reflect the diversity of human DC populations remains a major challenge in immunology (Luo & Dalod, 2020). MDDCs have been extensively used to study host-pathogen interactions because they are readily accessible, functionally relevant, and can be generated in large numbers. However, MDDCs represent an inflammatory DC population derived from circulating monocytes and do not fully recapitulate the phenotypic and functional diversity of conventional DC subsets found *in vivo*. Consequently, recent efforts have focused on generating DCs from HSPCs, thereby providing a more physiologically relevant experimental system (Lutz et al., 2023). Previous studies demonstrated the generation of *bona fide* cDC1, DC2 and pDCs populations from HSPC (Anselmi et al., 2020; Balan et al., 2018), whereas DC3-like populations have mainly been reported in more restricted differentiation settings (Bourdely et al., 2020). Based on the cell markers used in this study, we identified cDC1, DC2, DC3, moDCs and pDCs that were generated simultaneously from both BM- and CB-derived HSPCs. The lower pDC frequency observed in BM-derived cultures compared to CB-derived cultures likely reflects intrinsic differences in the starting progenitor pools. Cord blood is enriched for primitive cell progenitors, whereas adult bone marrow samples contain more heterogeneous populations and lineage-primed progenitors, which may reduce the capacity of cells to enter the pDC differentiation program under identical experimental conditions (Musumeci et al., 2019). These findings indicate that HSPC-derived cultures provide an experimentally tractable platform that reproduces key human DC subsets and can be used to investigate their interactions with HIV-1 under controlled conditions.

An important consideration when establishing an *in vitro* DC differentiation system is whether sufficient numbers of each subset can be generated to support robust and reproducible downstream analyses. In our cultures, approximately one-third of live cells acquired phenotypes consistent with differentiated DC populations, with DC2, DC3 and moDCs representing the predominant subsets. These proportions were broadly consistent with those reported in previous HSPC differentiation studies (Anselmi et al., 2020; Fiore et al., 2024). Notably, the relative abundance of most subset remained stable following IFN-α or LPS stimulation, except for DC3 cells, which exhibited a modest reduction under both activation conditions. This frequency change could be explained by cell death or by phenotypic shift due to changes in expression of lineage marker. Collectively, these findings indicate that immune activation does not substantially alter the overall representation of most DC subsets within the culture system and support the use of this platform for comparative analyses of DC subset function.

CD169 is a well-established interferon-inducible receptor involved in HIV-1 capture by macrophages and MDDC (Candor et al., 2024; Hammonds et al., 2017; Izquierdo-Useros et al., 2012; Pino et al., 2015; Puryear et al., 2013). In contrast, its expression in *bona fide* DC populations remains incompletely characterized (Rhodes et al., 2021). Previous studies using peripheral blood or tissue-derived samples have generally reported little or no detectable CD169 expression in cDC1, DC2, or pDC populations under steady-state conditions, although activation-associated transcriptional induction has been described (Rhodes et al., 2021; Winheim et al., 2025). In addition, circulating DC3 cells have been shown to upregulate CD169 under inflammatory conditions (Dutertre et al., 2019; Winheim et al., 2021). Consistent with these observations, our data demonstrate that both DC2 and DC3 subsets express detectable levels of CD169 that are markedly enhanced by IFN-α stimulation. Interestingly, BM- and CB-derived cultures exhibited distinct basal and inducible expression profiles, suggesting that developmental stage or progenitor composition may influence CD169 regulation (Da Silva et al., 2009). Although the biological basis of these differences remains to be determined, our findings identify DC2 and DC3 as interferon-responsive DC populations capable of expressing substantial levels of CD169.

HIV-1 capture by DC has been extensively documented in MDDC generated *in vitro* and in myeloid cells isolated from tissues (Akiyama et al., 2015; Izquierdo-Useros et al., 2012; Perez-Zsolt, Erkizia, et al., 2019; Pino et al., 2015; Puryear et al., 2013; Rhodes et al., 2021). However, much less is known about the capacity of conventional DC subsets to participate in this process. Previous studies suggested that DC2 cells can capture and transfer HIV-1 with limited efficiency (Parthasarathy et al., 2024; Rhodes et al., 2021; Ruffin et al., 2019; Silvin et al., 2017), whereas the role of DC3 in viral capture has been addressed recently, including reports describing the expression of classical and non-classical HIV-1 co-receptors (Buffa et al., 2025; Parthasarathy et al., 2024). Our results demonstrate that both DC2 and DC3 efficiently bind HIV-1-like particles and that viral capture is enhanced following IFN-α stimulation, closely paralleling CD169 expression levels. These findings expand current knowledge of HIV-1 interactions with myeloid DC populations and indicate that CD169-dependent viral capture is not restricted to monocyte-derived DCs. Consistent with previous observations, moDCs displayed the highest viral capture capacity and showed functional behavior similar to that of conventional MDDCs generated from peripheral blood monocytes.

Following HIV-1 capture, MDDCs accumulate viral particles within sac-like compartments through MEND-dependent mechanisms, a process that protects virions from degradation and facilitates *trans*- infection (Izquierdo-Useros et al., 2011, 2012; Laguía et al., 2025; Pino et al., 2015). Whether similar structures are formed by physiologically relevant DC populations has remained largely unexplored, although comparable virus-containing structures have been observed in CD11c^+^ cells within cervical tissue (Perez-Zsolt, Cantero-Pérez, et al., 2019). Here, we observed the formation of sac-like compartments in HSPC-derived DC cultures following VLP_HIV_ exposure, and the frequency of these structures increased after IFN-α stimulation, consistent with the enhanced viral capture observed under the same conditions. Although subset identity could not be resolved in the microscopy experiments, the presence of multiple *bona fide* DC populations within the cultures suggests that compartment formation is unlikely to be restricted exclusively to moDCs. Furthermore, pharmacological inhibition of MEND-associated pathways markedly reduced sac-like compartment formation, supporting the involvement of MEND-related mechanisms in HIV-1 internalization by HSPC-derived DCs and extending our previous observations in peripheral blood-derived MDDCs (Laguía et al., 2025). Finally, we demonstrate that HIV-1-loaded HSPC-derived DC cultures efficiently *trans*-infect CD4^+^ target cells and that this process is substantially reduced following CD169 blockade, as previously observed in MDDCs (Akiyama et al., 2015, 2017; Izquierdo-Useros et al., 2012; Pino et al., 2015; Puryear et al., 2013). These findings underscore the functional importance of CD169-mediated viral capture for downstream viral dissemination.

The present study has several limitations that should be considered. Our study relies on a flow cytometry panel based on previously described surface markers to approximate subset identity but lacks RNA-seq profiling to rigorously define and distinguish all DC subsets generated from CD34^+^ HSPCs. Confocal microscopy and *trans*-infection experiments were performed on mixed HSPC-derived cultures; therefore sac-like compartment formation and viral transfer to target cells could not be directly assigned to individual DC subsets. Although CD169 blockade significantly reduced *trans*-infection, residual viral transfer was still detected under all conditions. Therefore, the present study does not exclude the participation of additional uptake or transfer pathways acting in parallel with CD169. On the other hand, our spectral cytometry panel did not include certain surface markers, such as CD163, Axl, and Siglec-6, that are used to identify certain DC populations that were not considered to evaluate their potential contribution to HIV-1 capture. Future studies combining subset purification with imaging and functional analyses will be required to define the relative contribution of individual DC populations to CD169-dependent viral capture, intracellular trafficking, and viral dissemination.

In conclusion, human CD34^+^ HSPC-derived cultures provide a versatile and physiologically relevant platform for investigating HIV-1 interactions across multiple dendritic cell populations. Collectively, our findings demonstrate that CD169-dependent HIV-1 capture is not restricted to monocyte-derived DCs but is also a property of conventional DC subsets, particularly DC2 and DC3. Although MDDCs remain a valuable experimental model for studying CD169 biology and virus-host interactions, our results broaden the spectrum of DC populations potentially involved in HIV-1 capture. Our study establishes HSPC-derived cultures as a useful system for dissecting the contribution of distinct myeloid cell subsets to viral dissemination.

## Abbreviations

DC: Dendritic Cells
BM: Bone marrow
CB: Cord blood
MDDC: Monocyte-derived dendritic cells
MEND: Massive Endocytosis
HSPC: Hematopoietic stem and progenitor cells

## Acknowledgements

We thank the Scientific and technical services (IrsiCaixa) for the blood sample processing and PBMCs isolation. We are grateful to J. Puñet-Ortiz and S. Monreal from IGTP Cytometry Core Facility for their technical contribution. Some figures were created with BioRender.com.

AI Use Declaration: Artificial intelligence–assisted tools (Microsoft Copilot/GPT-based language models) were used to support grammar refinement and text structuring during manuscript preparation. All scientific content, data interpretation, and final text were generated solely by the authors that take full responsibility for the accuracy and integrity of the manuscript.

## Author contributions

FL: Conceptualization, Data Curation, Formal Analysis, Investigation, Methodology, Validation, Visualization, Writing – original draft, Writing – review & editing

PP: Investigation, Methodology, Writing – review & editing

MAV: Investigation, Methodology, Writing – review & editing

JIP: Investigation, Methodology, Writing – review & editing

CMP: Investigation, Methodology, Writing – review & editing

JMP: Conceptualization, Funding acquisition, Investigation, Project Administration, Supervision, Validation, Writing – original draft, Writing – review & editing

PRI: Conceptualization, Data Curation, Formal Analysis, Investigation, Methodology, Project Administration, Supervision, Validation, Visualization, Writing – original draft, Writing – review & editing

## Figure and tables Captions

**Figure S1.**
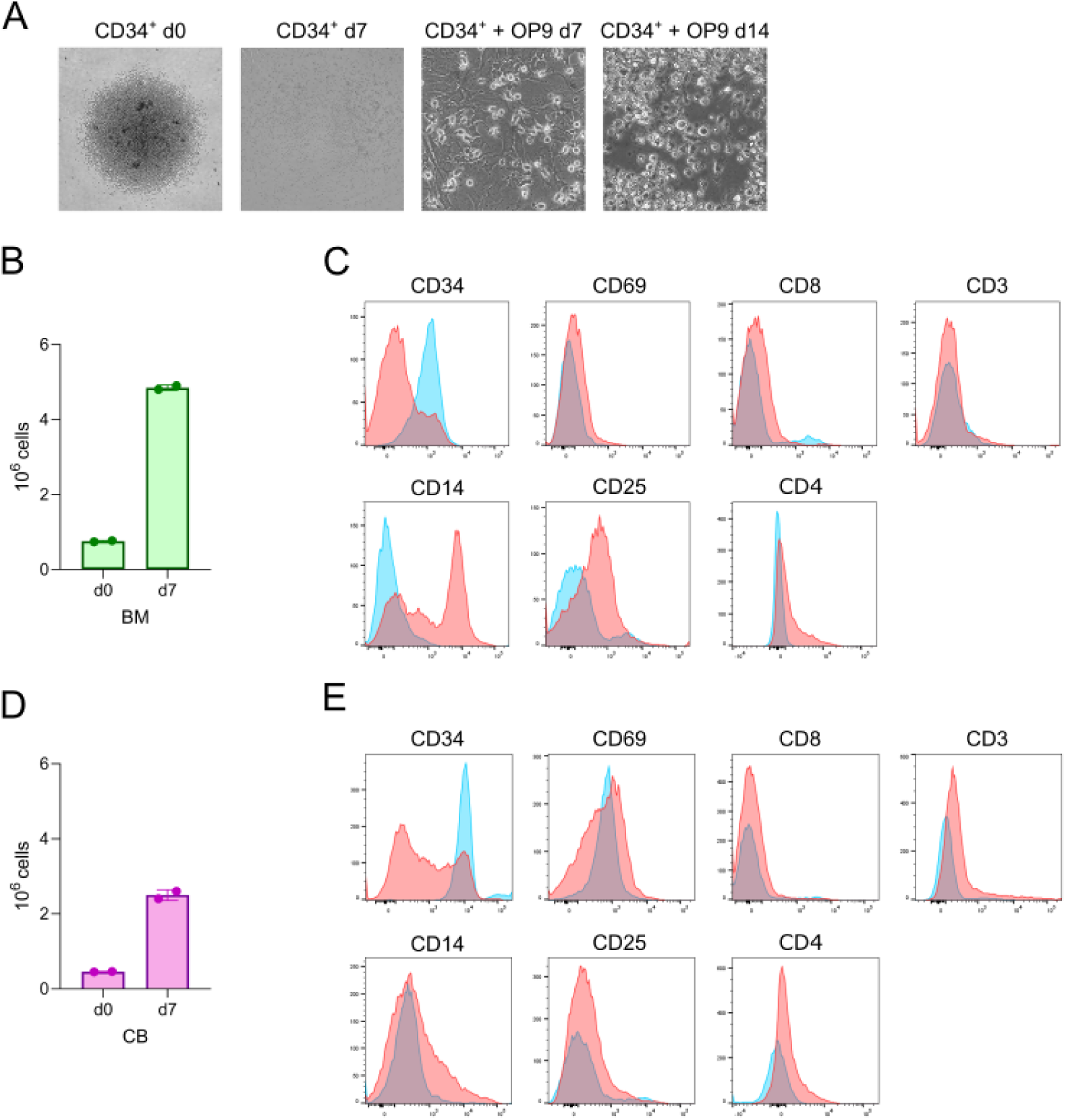
CD34^+^ HSPCs expand and initiate differentiation during the expansion phase. HSPCs were cultured under expansion conditions for 7 days and analyzed for changes in cell number and immunophenotype. **(A)** Representative wide-field microscopy images of HSPCs at day 0 and day 7 of expansion and during co-culture with OP9 feeder cells. **(B)** Quantification of BM-derived HSPC expansion between day 0 and day 7. **(C)** Flow cytometry histograms showing expression of the indicated markers in BM-derived HSPCs at day 0 (blue) and day 7 (red). **(D)** Quantification of CB-derived HSPC expansion between day 0 and day 7. **(E)** Flow cytometry histograms showing expression of the indicated markers in CB-derived HSPCs at day 0 (blue) and day 7 (red).

**Figure S2.**
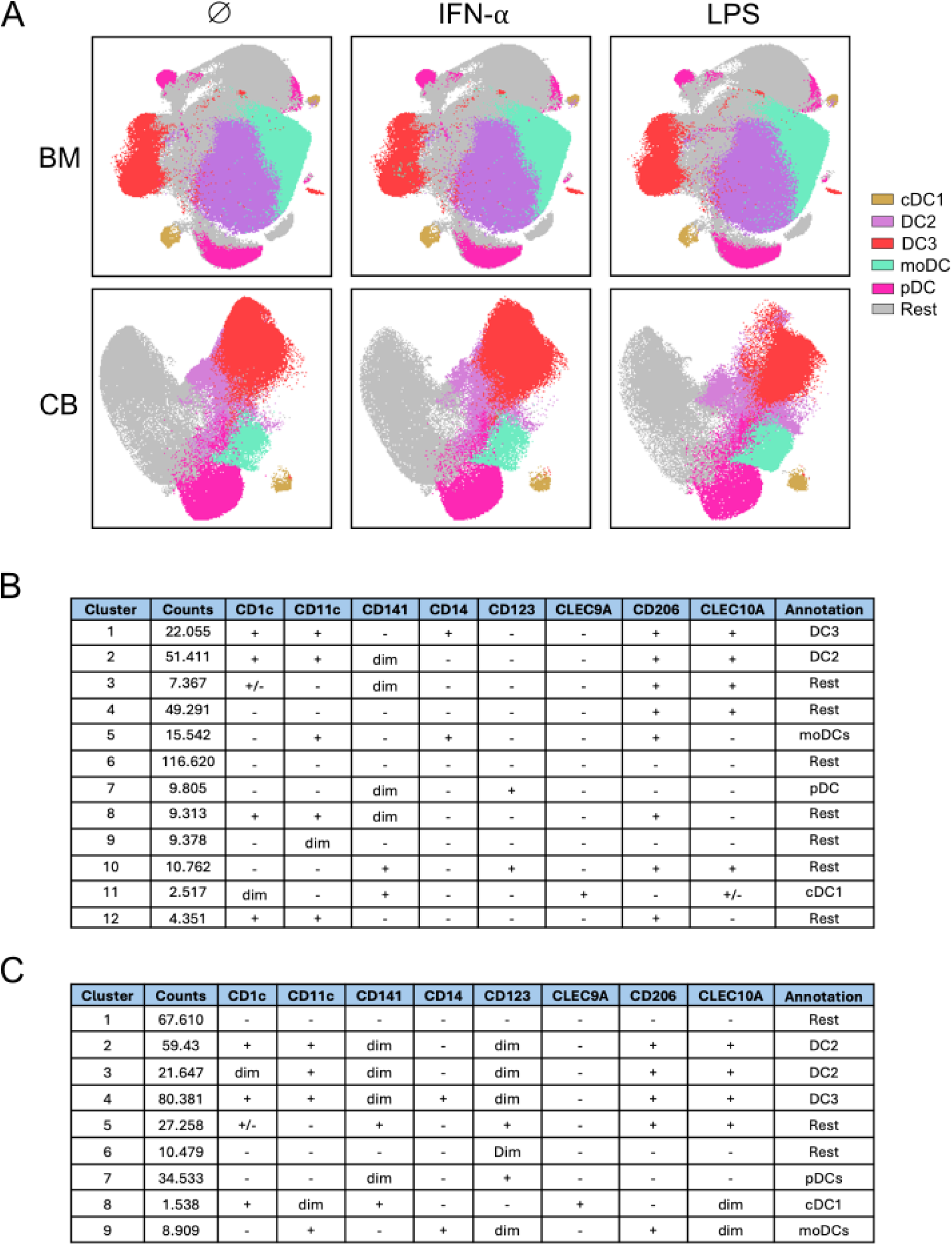
FlowSOM analysis validates the identification of DC subsets by manual gating generated from CD34^+^ HSPCs. **(A)** Uniform manifold approximation and projection for dimension reduction (UMAP) representation of FlowSOM clusters obtained from differentiated BM- and CB-derived cultures representing the five major DC subsets. Clusters were defined using the same marker panel used for the manual gating strategy. Subsets whose markers did not match with described populations were grouped as rest. One representative experiment is shown for each type of culture. **(B)** Cell count, marker annotation and DC-cell population in BM-derived DCs. **(C)** Cell count, marker annotation and DC-cell population in CB-derived DCs.

**Figure S3.**
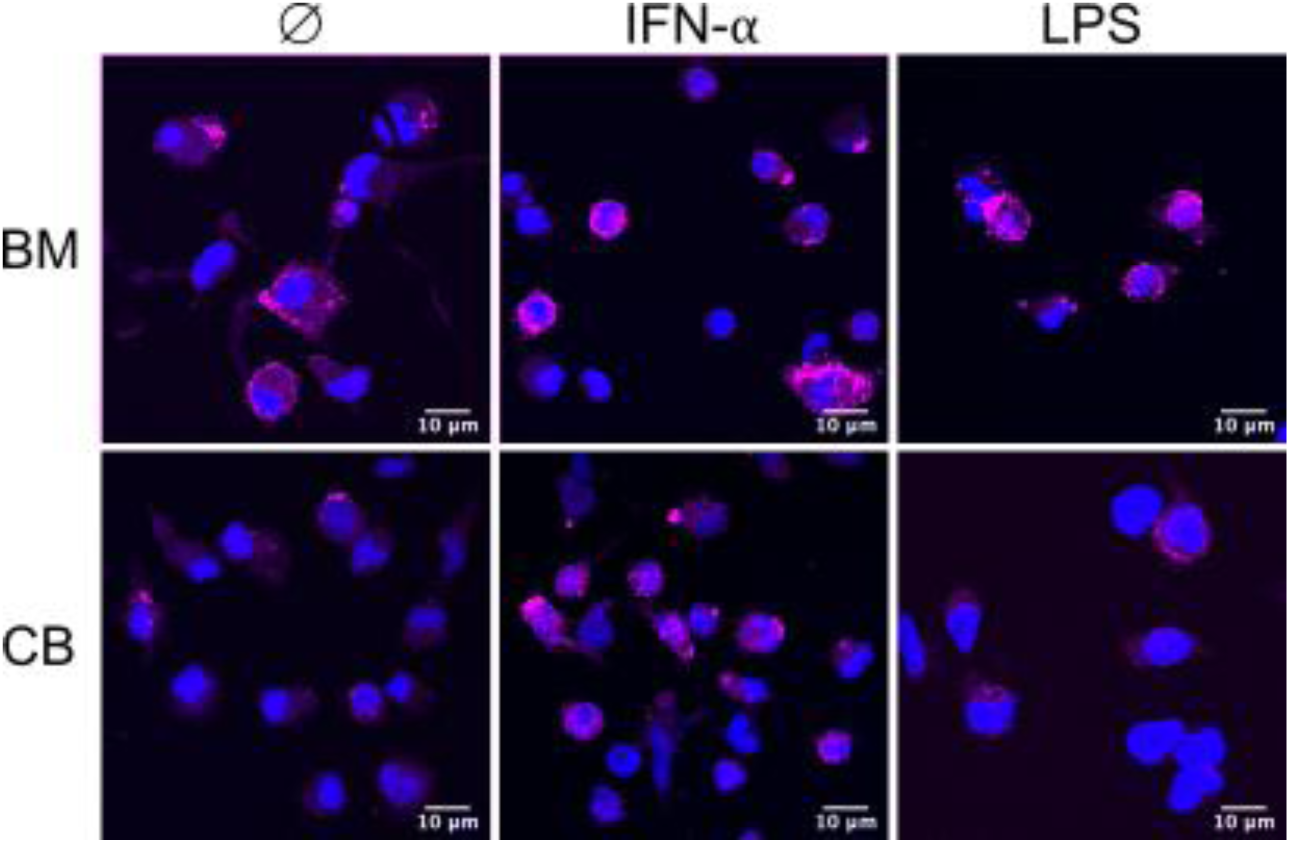
Confocal microscopy analysis of CD169 expression in HSPC-derived DCs. Representative confocal images of BM- and CB-derived cultures under control, IFN-α, and LPS conditions. Cells were stained with an anti-CD169 monoclonal antibody. Images represent projections of 5–10 optical sections spanning the equatorial region of the cells.

**Figure S4.**
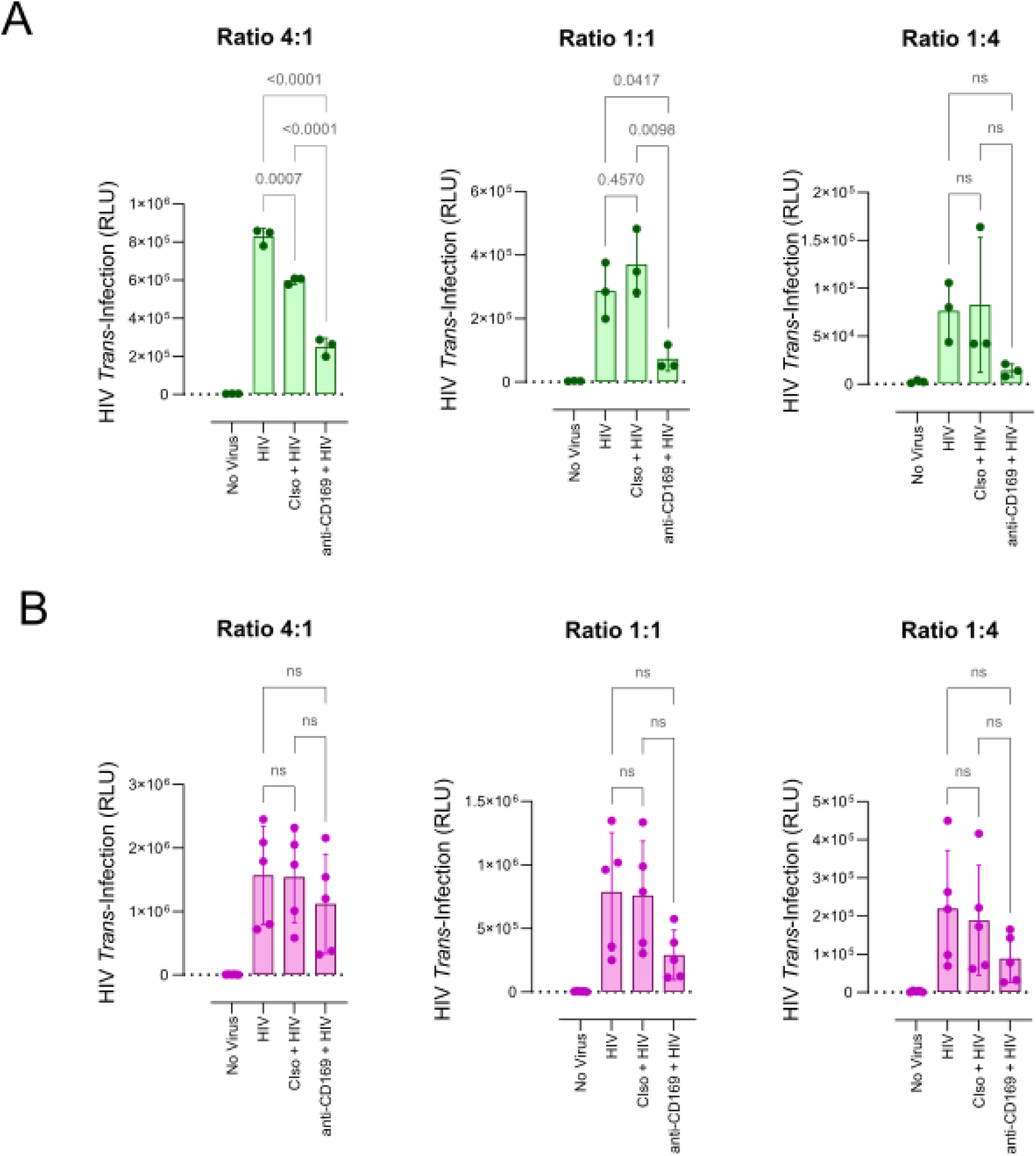
Absolute HIV-1 *trans*-infection values in HSPC-derived DC cultures. The experimental conditions correspond to those shown in Figure 5. Bars represent mean values, and error bars indicate the standard deviation. Statistical significance was assessed by one-way ANOVA. **(A-B)** Absolute luciferase activity expressed as relative light units (RLU), measured in TZM-bl cells following co-culture with BM-derived **(A)** and CB-derived **(B)** DCs at DC:TZM-bl ratios of 4:1, 1:1 and 1:4.

